# Adenosine 5’-triphosphate (ATP) forms protein-free and responsive condensates in crowded environments

**DOI:** 10.64898/2026.03.22.713448

**Authors:** Yuchao Wang, Feipeng Chen, Parrik Dang Kow, Ho Cheung Shum

## Abstract

Adenosine 5’-triphosphate (ATP) is found to form biomolecular condensates with proteins. However, without complementary proteins, the small size and high charge density of ATP molecules create substantial electrostatic and entropic barriers that prevent them from forming condensates. Here, we find that macromolecular crowding overcomes these energetic barriers, promoting ATP molecules to self-associate and form protein-free liquid-like condensates through screened electrostatic repulsion and enhanced hydrogen bonding. Importantly, ATP condensates are responsive to multiple stimuli and create distinct microenvironments that selectively enrich various guest molecules and protect ribonucleic acids from DNAzyme cleavage. These findings uncover important roles of ATP in forming dynamic, chemically distinct condensates via homotypic interactions, potentially expanding its classical view beyond a canonical energy carrier to a structural and regulatory architect in cellular physiology and prebiotic chemistry.

## Main Text

Adenosine 5’-triphosphate (ATP) serves as the primary energy currency to fuel numerous biochemical reactions and power cellular activities. Beyond this fundamental role, ATP also participates in the formation of membraneless biomolecular condensates (*1-3*). Despite lacking physical membranes, these biomolecular condensates create physically separated and chemically unique compartments that orchestrate diverse biological reactions in cells (*4-7*) and feature as promising protocells to evolve under prebiotic conditions (*8-10*). Biomolecular condensates are generally formed by liquid-liquid phase separation (LLPS) of macromolecules, such as proteins and nucleic acids, that interact through associative multivalent interactions (*11-14*). These interactions reduce electrostatic repulsion and minimize translational entropy among macromolecules, resulting in a thermodynamic landscape conducive to LLPS (*15, 16*). As a result, previous studies have found that negatively charged ATP interacts with many intrinsically disordered proteins with positive residues to form complex condensates (*17-19*).

However, without binding partners, ATP molecules are thermodynamically unfavorable for forming condensates by themselves. As a highly charged small molecule, ATP has inherent high charge density, creating significant electrostatic barriers for its self-interaction into condensates (Fig. 1A, i) (*1, 20*). In addition, the small size of ATP further confers high diffusivity and consequently high translational entropy, imposing large entropic barriers to its condensation (Fig. 1A, ii) (*21*). Furthermore, previous studies have shown that ATP functions as a hydrotrope, solubilizing aggregation-prone protein condensates potentially by binding to and sequestering proteins (Fig. 1A, iii) (*22, 23*). These intrinsic characteristics collectively suggest that ATP molecules are energetically unfavorable for forming condensates by themselves via homotypic interactions.

**Fig. 1.**
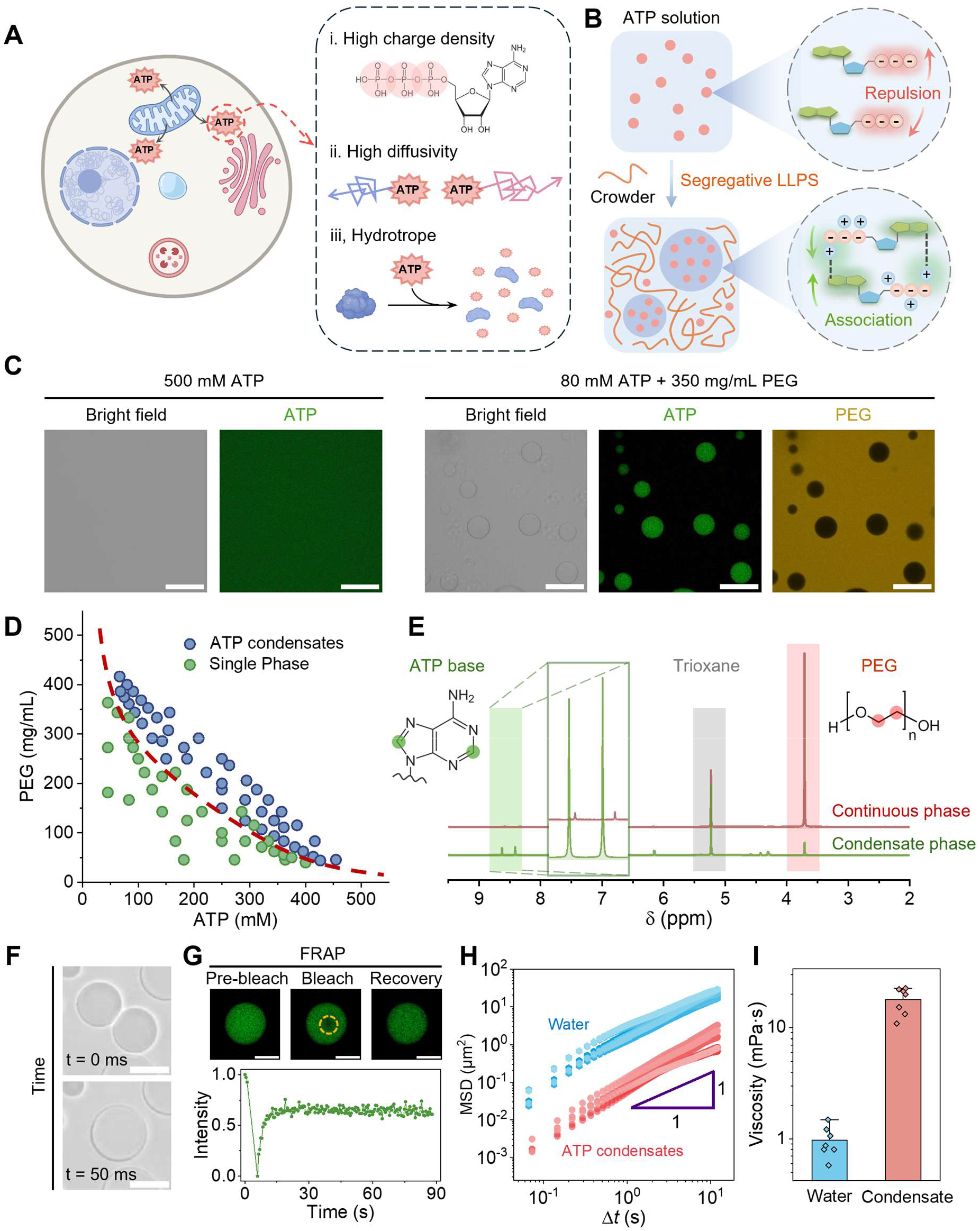
ATP molecules form liquid-like condensates in the presence of crowders. (**A**) Schematic illustration of the physicochemical characteristic of ATP molecules, highlighting their charged triphosphate groups, high diffusivity, and hydrotropic properties. (**B**) Schematic illustration showing ATP condensation from a homogeneous ATP solution upon addition of crowder, mediated by a crowding-induced interaction switch among ATP molecules. (**C**) Representative images of ATP solutions with and without crowders. The introduction of PEG as crowders lead to the formation of ATP condensates (*c*_PEG_ = 350 mg/mL and *c*_ATP_ = 80 mM). Fluorescein isothiocyanate (FITC) labelled Dextran and Rhodamine (Rb) labelled PEG are utilized to track ATP and PEG within condensates and the continuous phase. Scale bars are 100 μm. (**D**) Phase diagram of ATP-PEG mixture, delineating the homogeneous single-phase regime (green circles) and the ATP condensate regime (blue circles) as functions of PEG and ATP concentrations. The red dashed curve indicates the binodal curve between two regimes. (**E**) ^1^H NMR spectrum of the condensate (green) and continuous phases (red). Trioxane is used as a calibration molecule. (**F**) Representative images of rapid coalescence between two ATP droplet condensates. Scale bars are 10 μm. (**G**) Fluorescence recovery after photobleaching (FRAP) analysis of ATP condensates. Fluorescence images and a corresponding plot of fluorescence intensity show rapid recovery over time in a photobleached region (the dashed yellow circle). Scale bars are 10 μm. (**H**) Mean squared displacement (MSD) versus lag time *Δt* of fluorescent microparticles embedded in water (blue hexagons) and ATP condensates (red circles), *n* = 7. (**I**) Comparison of calculated viscosity between water and ATP condensates, *n* = 7.

In this work, we surprisingly find that ATP form droplet condensates via homotypic interactions in the presence of macromolecular crowders. This condensation process is driven by a synergistic interplay of screened electrostatic repulsion and enhanced hydrogen bonding among ATP molecules, leading to segregative LLPS with crowders. Mediated by these weak homotypic interactions, ATP condensates exhibit liquid-like behaviors, rapidly merging upon contact and recovering the fluorescence intensity after photobleaching. In contrast to condensates formed by associative LLPS, ATP condensates dynamically form and disassemble in response to environmental stimuli, such as temperature, pH, and concentration. Consequently, they act as dynamic hubs to selectively concentrate and release diverse biomolecules. More importantly, ATP condensates are found to sequester ribonucleic acids (RNA) and protect them from being cleaved. These findings reveal an important yet previously unrecognized role of ATP in forming liquid-like condensates via homotypic interactions, potentially expanding its canonical function as the cellular energy currency.

### ATP forms liquid-like droplet condensates in crowded environments via segregative phase separation

As a highly charged small molecule, ATP is not expected to form condensates through homotypic interactions due to its high charge density, high translational entropy, and hydrotropic property (Fig. 1, A and B). Therefore, ATP solutions remain homogeneous and transparent even at very high concentrations up to 500 mM (Fig. 1, B and C). Surprisingly, upon adding polyethylene glycol (PEG) as macromolecular crowders, droplet condensates spontaneously appear in the solution (Fig. 1, B and C). Fluorescence imaging reveals that ATP is concentrated within the droplet condensates, while PEG is enriched in the surrounding continuous phase (Fig. 1C), suggesting a segregative LLPS between ATP and PEG. By measuring the phase diagram of ATP-PEG mixture, we find that condensates form when ATP and PEG concentrations exceed critical thresholds. These critical points delineate a binodal curve, above which ATP condensates form (blue circles), whereas below which there are no condensates (green circles) (Fig. 1D). Notably, the critical ATP concentration required for phase separation decreases with increasing PEG concentration and vice versa (Fig. 1D). Consistent with the fluorescence imaging, this inverse relationship suggests a segregative LLPS mechanism, by which ATP and PEG demix with each other to form distinct phases (*20, 24, 25*).

To further confirm the segregative LLPS, we detect the composition of the condensate and the continuous phase using ^1^H nuclear magnetic resonance (NMR). Characteristic peaks at ∼8.5 ppm and ∼3.7 ppm are first identified from ATP base protons and PEG backbone protons respectively (fig. S1). NMR spectrum of the continuous phase reveals a strong peak at ∼3.7 ppm from PEG backbone protons, while the peak at ∼8.5 ppm from ATP base protons is negligible (Fig. 1E). In contrast, the condensate phase exhibits a strong peak from ATP base protons but a weak peak from PEG backbone protons (Fig. 1E). These results confirm that PEG is mostly enriched in the continuous phase while ATP is concentrated within the condensate phase. Moreover, mixing the continuous phase with dextran (DEX) leads to demixing of DEX into droplets (fig. S2), confirming that PEG concentration is sufficiently high in triggering the segregative LLPS with DEX (*26, 27*). Conversely, the condensate phase remains homogeneous when mixing with DEX but readily forms condensates upon introducing poly-l-lysine (PLL) even at very low concentrations (figs. S2 and S3), suggesting the locally enriched ATP inside condensates. Taken together, these results demonstrate that the macromolecular crowder PEG can overcome ATP’s entropic and electrostatic barriers, driving its condensation through a segregative LLPS.

We further find that these condensates exhibit liquid-like properties. For example, they quickly merge into larger spherical ones upon contact (Fig. 1F). Fluorescence recovery after photobleaching (FRAP) experiments also show that a bleached area inside the condensates recovers rapidly (Fig. 1G). To quantitatively probe their properties, we track fluorescent microparticles that preferentially partition into ATP condensates (fig. S4). Single-particle trajectory analysis shows that microparticles move randomly inside ATP condensates and bulk water (fig. S5). Fitting the mean squared displacement (MSD) of microparticles with MSD(Δ*t*) = 4*D*Δ*t*^*α*^, we obtain the scaling exponent *α* and the diffusion coefficient *D*. The MSD of microparticles in both water and ATP condensate exhibits classic Brownian motions with *α*≈1 (Fig. 1H). Differently, the diffusion coefficient D of microparticles within ATP condensates is about 0.026 μm^2^/s, which is slower than that of 0.487 μm^2^/s in water. Applying the Stokes-Einstein equation 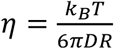 (where *k*_*B*_ is the Boltzmann constant, *T* is the absolute temperature, and *R* is the particle radius), the viscosity of ATP condensates is approximately 17.90 mPa·s, one order of magnitude higher than that of water (Fig. 1I). Nonetheless, the viscosity of ATP condensates is significantly lower than that of many condensates formed via associative LLPS (10^3^-10^5^ mPa·s), underscoring their highly liquid-like nature (*28-31*).

### Screened electrostatic repulsion and enhanced hydrogen bonding lead to ATP condensation

To elucidate the condensation mechanism, we systematically screen the interactions involved in ATP condensates by introducing small molecular perturbants (*32, 33*). We start with a condition in the single-phase regime (*c*_ATP_ = 60 mM, *c*_PEG_ = 300 mg/mL) where condensates could not form. By titrating different kinds of salts into the system, condensates are formed above critical salt concentrations. We observe that all kinds of salts (NaCl, KCl, MgCl_2_, CaCl_2_) promote ATP condensation (Fig. 2, A and B, and fig. S6). In particular, divalent cations (Ca^2+^, Mg^2+^) are more effective than monovalent cations (Na^+^, K^+^), following a trend of Mg^2+^ ≈ Ca^2+^ > Na^+^ > K^+^ (Fig. 2, B and C, and fig. S7). This trend aligns with the Hofmeister series that marks the ionic capacity to screen electrostatic interactions (*32, 34, 35*). Therefore, these results suggest that ATP condensation is facilitated by screened electrostatic repulsion. This is further supported by enhanced ATP condensation in the presence of alcohols, such as ethanol and 1,6-hexanediol (1,6-HD) (Fig. 2, A and B). Alcohols lower dielectric constants of co-solvents and thus increase the binding energy between ATP and counterions (*36, 37*). As a result, ATP molecules are neutralized with reduced release of counterions and hydrogen ions, diminishing the electrostatic repulsion among themselves. Consequently, adding salts and alcohols shifts the binodal curves to the bottom left, further lowering the minimal concentrations of ATP and PEG required for phase separation (Fig. 2D). In addition to electrostatic interactions, ATP molecules potentially interact by forming hydrogen bonds (*38-40*). We observe that ATP condensates gradually dissolve with the addition of urea, a potent hydrogen-bond disruptor (Fig. 2, E and F), suggesting the critical role of hydrogen bonding in stabilizing ATP condensates (*33*). In particular, adding urea inhibits ATP condensation and shifts the binodal curves to the upper right (fig. S8). Combined with that ATP condensates could form in the presence of PEG without additional salts, alcohols and urea, it implies that crowding conditions alone could suppress counterion release from ATP molecules, thereby weakening their electrostatic repulsion, and promote hydrogen bonding between ATP molecules, thus leading to enhanced self-associations (*41*).

**Fig. 2.**
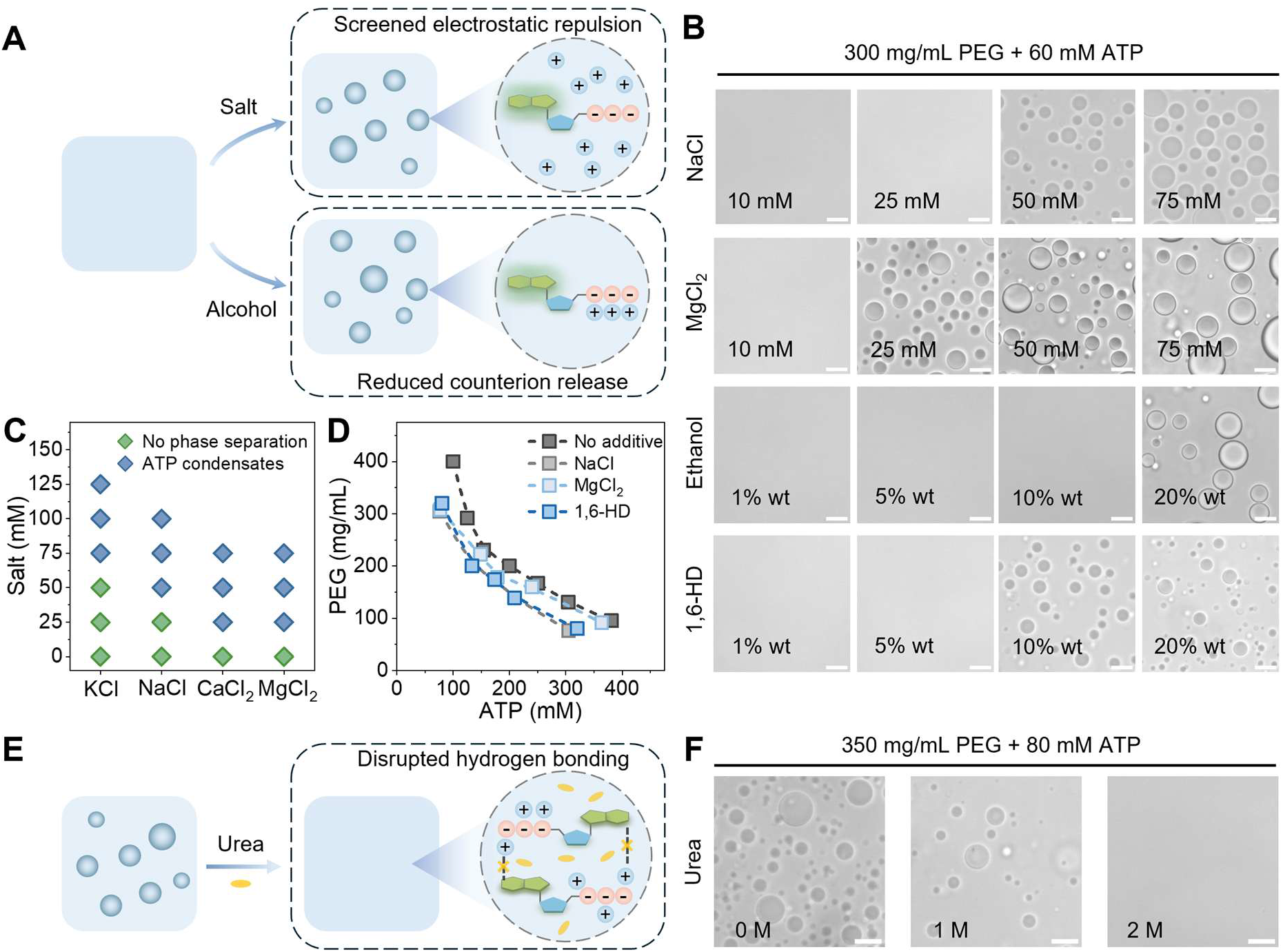
Electrostatic screening and hydrogen bonding lead to ATP condensation. (**A**) Schematic illustration of the impact of salt and alcohol on ATP condensation. The addition of salt and alcohol promotes ATP condensation at lower ATP and PEG concentrations. (**B**) Representative images show ATP solutions (*c*_PEG_ = 300 mg/mL, *c*_ATP_ = 60 mM) with different concentrations of salts and alcohols. Scale bars are 10 μm. (**C**) Quantification of critical salt concentrations for salt-promoted ATP condensation (*c*_PEG_ = 300 mg/mL, *c*_ATP_ = 60 mM). (**D**) Phase diagrams and binodal curves of ATP™PEG mixture with different additives: 100 mM NaCl, 20 mM MgCl_2_, 10 wt% 1,6-HD. (**E**) Schematic illustration of the impact of urea on ATP condensation. The addition of hydrogen bonding disruptor urea inhibits ATP condensation. (**F**) Representative images show ATP condensates (*c*_PEG_ = 350 mg/mL, *c*_ATP_ = 80 mM) with different concentrations of urea. Scale bars are 10 μm.

Notably, in the absence of PEG, ATP solutions remain homogeneous even at high concentrations of salts or alcohols (figs. S9 and S10), highlighting the indispensable role of crowding in driving ATP condensation. To explore how crowders modulate ATP condensation, we further examine the effects of PEG chain length. Mean-field theory suggests that PEG with higher chain length is more effective in promoting the segregative phase separation (Fig. 3A), potentially caused by enhanced excluded volume effects and reduced translational entropy of polymers with longer chains (*14, 42*). Indeed, turbidity measurements reveal that PEG 2K or 8K, is more efficient than its low-molecular-weight counterparts PEG 600 in inducing ATP condensation (Fig. 3B). For example, at a fixed ATP concentration, a notable ∼50% turbidity increase is triggered by adding PEG 8K at 250 mg/mL, whereas it requires a concentration of 350 mg/mL for PEG 600 (Fig. 3B). The monomer ethylene glycol (EG) even fails to induce condensate formation at all tested concentrations up to 500 mg/mL. As a result, introducing PEG of higher molecular weights shifts the binodal curve more toward the bottom left (Fig. 3C), effectively lowering the critical concentration required for phase separation. To test the universality of crowding effects, we substitute PEG with alternative crowding agents, including poly(2-ethyl-2-oxazoline) (PEtOx), dextran, and Ficoll. PEtOx induces ATP condensation above critical concentration thresholds in a manner similar to PEG (figs. S11 and S12). However, ATP can not form condensates by adding dextran and Ficoll across a wide range of concentrations (figs. S11 and S12). The distinct effects of different crowders imply that ATP condensation is not simply driven by crowding-induced excluded volume effects. Since PEG and PEtOx are considered more hydrophobic than dextran and Ficoll (*41*), it suggests that hydrophobic crowders better enhance self-associations among ATP molecules.

**Fig. 3.**
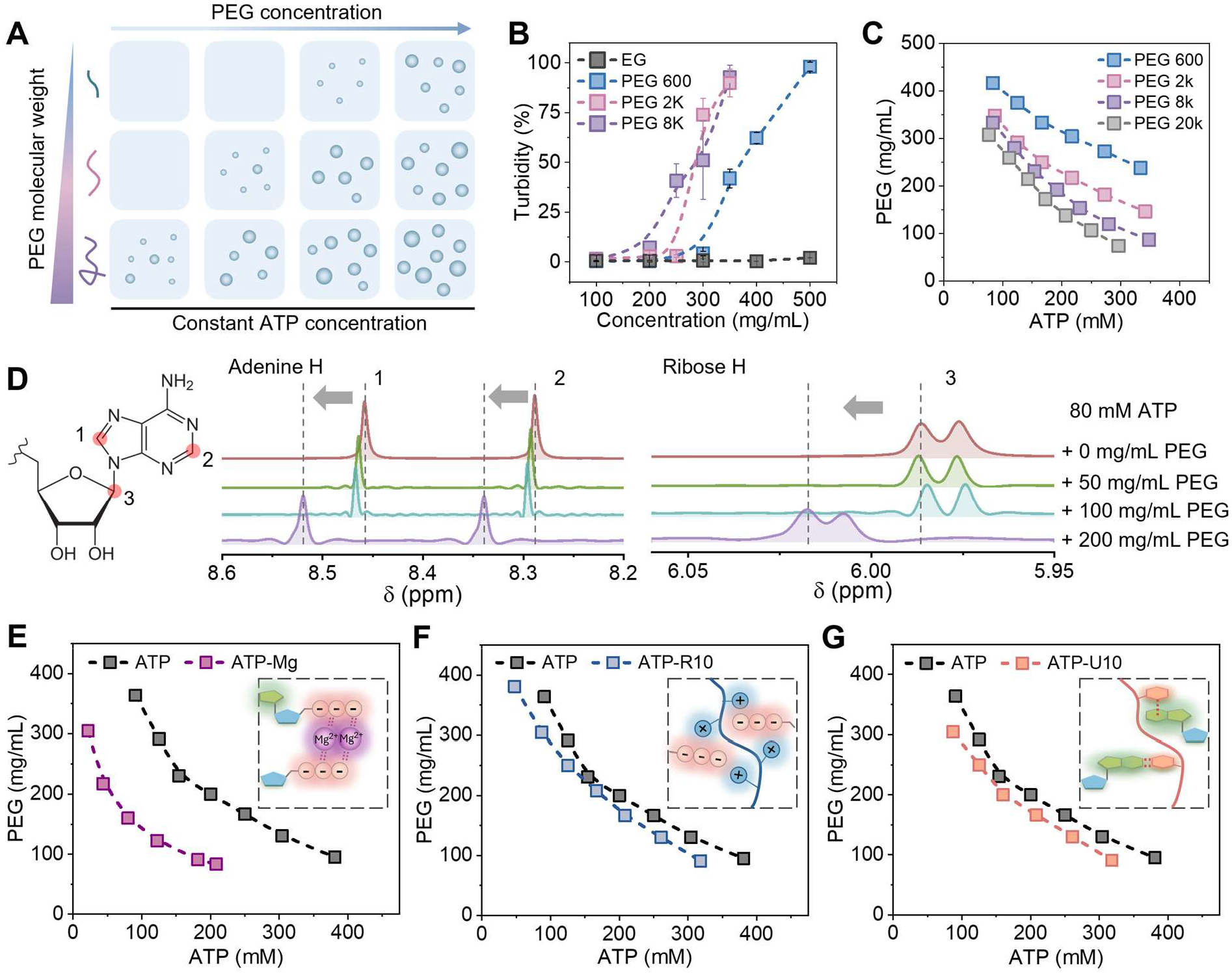
Modulation of ATP condensation by crowders and physiologically relevant molecules. ((**A**) Schematic illustration of the impact of PEG polymerization on ATP condensation. PEG with higher molecular weight is more effective at inducing ATP condensation. (**B**) Turbidity of ATP solutions measured at OD 500 nm in the presence of PEG with different molecular weight and concentrations. *c*_ATP_ = 150 mM. (**C**) Phase diagrams and binodal curves of ATP™PEG mixture with PEG of varying molecular weights. (**D**) Schematic of highlighted ATP adenine base and ribose sugar protons and a typical ^1^H NMR spectrum acquired with 80 mM ATP and increasing PEG concentrations (from top to bottom). (**E**) Phase diagram and binodal curve of ATP™PEG mixture containing Mg^2+^ added at equimolar concentration to ATP. (**F**) Phase diagram and binodal curve of ATP™PEG mixture with the addition of R10 added at molar concentration of 10000:1 (ATP: R10). (**G**) Phase diagram and binodal curve of ATP™PEG mixture with the addition of U10 added at molar concentration of 400:1 (ATP: U10).

To further confirm the potential interactions underpinning ATP condensation, we apply ^1^H NMR spectroscopy to monitor the proton chemical shifts of ATP functional motifs in the presence of crowders (*39, 43*). Intriguingly, increasing PEG concentration induces consistent downfield shifts of the adenine base and ribose sugar protons (Fig. 3D). This indicates the increased deshielding of protons, potentially caused by enhanced hydrogen bonding (*44*). Combined with urea-inhibited ATP condensation, the observed downfield shift indicates that the introduction of PEG enhances hydrogen bonding between ATP molecules. By contrast, the addition of dextran induces opposite upfield shifts of protons on the adenine base and ribose sugar (fig. S13). Similar upfield shifts are observed when ATP concentration increases (fig. S14). These upfield shifts are potentially caused by enhanced aromatic ring current effects and the consequent magnetic shielding at increased ATP concentrations (*45-47*). Together with previous small molecule perturbation experiments, these results collectively demonstrate that ATP condensation in crowded environments is driven by a synergy of screened electrostatic repulsion and enhanced hydrogen bonding among ATP molecules.

With this understanding, we further demonstrate the promotion of ATP condensation by introducing physiologically relevant components (Fig. 3, E to G). In cells, most of ATP are complexed with magnesium ions (Mg^2+^) (*48, 49*). Interestingly, we find the addition of Mg^2+^ at equimolar concentration to ATP remarkably promote ATP condensation. In particular, the addition of Mg^2+^ significantly shifts the binodal curve to lower ATP and PEG concentrations (Fig. 3E). This promoted condensation is consistent with the role of Mg^2+^ in screening electrostatic repulsions and potentially bridging ATP molecules (*49, 50*). In addition, oligopeptides and oligonucleotides, such as decapeptides of arginine (R10) and oligo(uridylic acid) RNA (U10), also slightly promote the ATP condensation and shift the binodal curve to the bottom left (Fig. 3, F and G). Therefore, although the required ATP and PEG concentrations for segregative LLPS are not physiologically relevant, additional associative agents that potentially exist in cells could effectively lower the threshold concentrations. In these cases, the segregative LLPS of ATP is coupled with associative phase transitions.

### Environment-responsive ATP condensate dynamics

Typical ATP-polycation condensates are formed by strong heterotypic interactions and remain stable in response to environmental fluctuations (*20, 51*). For example, ATP-PLL condensates remain very stable across a wide range of variations in concentration, pH, or temperature (figs. S15 to S18). They even form solid-like aggregates under basic conditions (fig. S18). However, as ATP condensates are driven by weak homotypic interactions, we find they are very responsive to environmental changes, dynamically assembling and dissolving in response to stimuli such as temperature, pH, and concentration fluctuations (Fig. 4A). For example, using a concentration condition in the single-phase regime, ATP condensates remain dissolved at 20 °C but form at 60 °C (Fig. 4B). Subsequent cooling to 20 °C again leads to a reversible dissolution of ATP condensates (Fig. 4B). In addition, raising the temperature from 20 °C to 60 °C shifts binodal curves to the bottom left (Fig. 4C and fig. S19). These results suggest that ATP condensates exhibit a lower critical solution temperature (LCST) behavior.

**Fig. 4.**
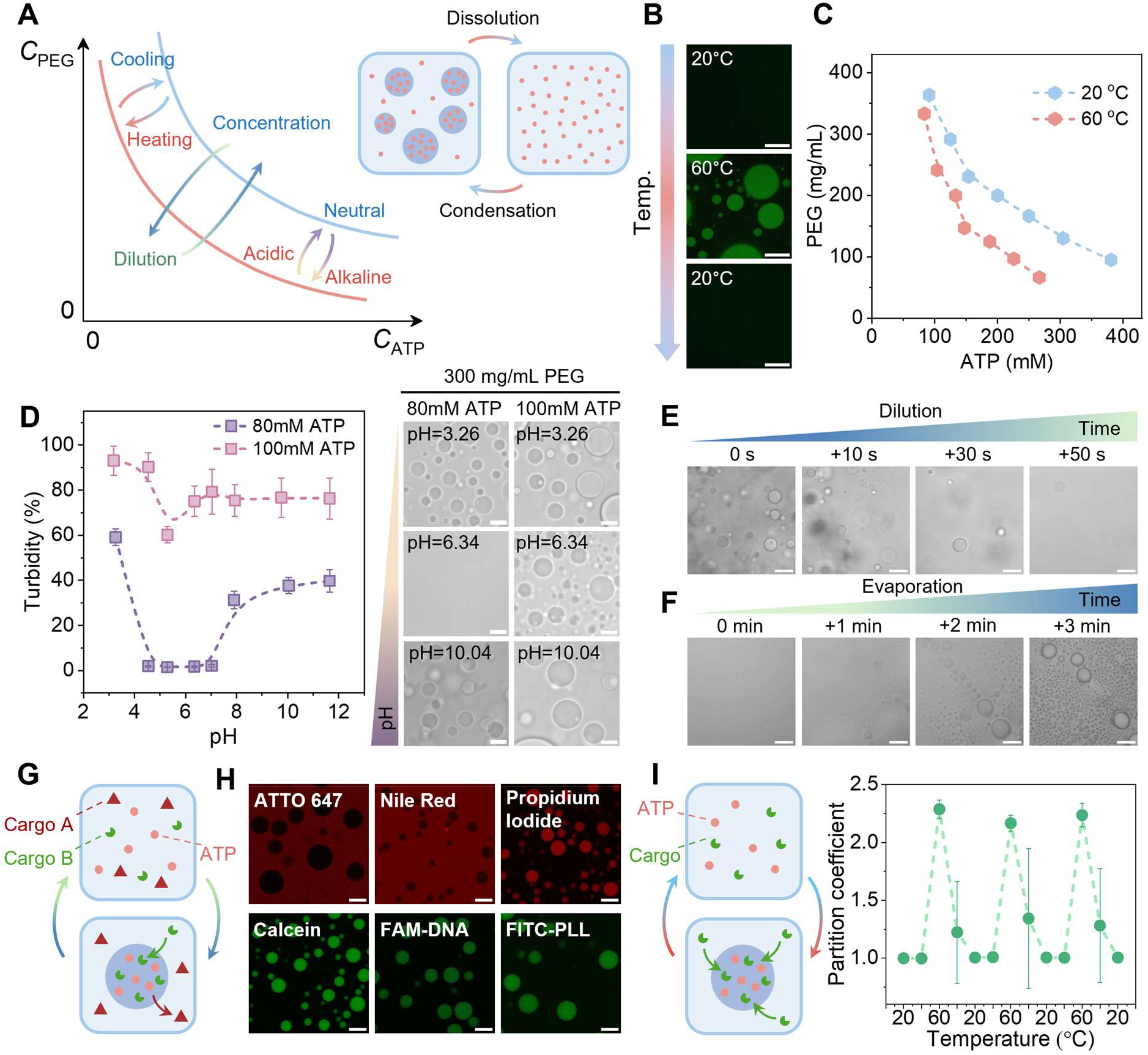
Environment-responsive ATP condensate dynamics. (**A**) Schematic illustration of dynamic formation and dissolution of ATP condensates in response to changes in temperature, pH, and concentration. (**B**) Representative fluorescence images of ATP condensates (*c*_PEG_ = 175 mg/mL and *c*_ATP_ = 175 mM) undergo a heating and cooling thermal cycle. Condensates exhibit a lower critical solution temperature (LCST) behavior. Calcein is used to visualize the ATP condensates. Scale bars are 50 μm. (**C**) Phase diagrams and binodal curves of ATP condensates at 20 °C and 60 °C. (**D**) Turbidity and representative optical images of ATP-PEG solutions (*c*_ATP_ = 80 mM and *c*_ATP_ = 100 mM) with constant PEG concentration of 350 mg/mL under different pH conditions, *n* = 5. Scale bars are 10 μm. (**E**)(**F**) Representative images of the dissolution of ATP condensates (*c*_PEG_ = 350 mg/mL and *c*_ATP_ = 80 mM) upon twofold dilution with water and their subsequent reformation via water evaporation induced by heating at 50 °C. Scale bars are 100 μm. (**G**) Schematic illustration showing the selective partitioning of guest molecules within ATP condensates. (**H**) Representative fluorescence images showing the selective partitioning of diverse guest molecules in ATP condensates (*c*_PEG_ = 350 mg/mL and *c*_ATP_ = 80 mM) or the continuous phase. Scale bars are 20 μm. (**I**) Schematic illustration of cargo release and enrichment controlled by the dissolution and formation of condensates and evolution of the partition coefficient of calcein (0.25mg/mL) in ATP condensates (*c*_PEG_ = 175 mg/mL and *c*_ATP_ = 175 mM) throughout three cycles of heating and cooling, measured by confocal microscopy. Data is represented as mean ± standard deviations, *n* = 20.

In addition, we find that ATP condensates exhibit a reentrant phase-separation behavior in response to pH. As pH increases from acidic to alkaline titrated by NaOH, the solution turbidity undergoes gradual decrease followed by recovery. (Fig. 4D and fig. S20). The decreased turbidity from basic to neutral pH indicates the dissolution of condensates which is likely caused by ATP deprotonation (*52*). At low pH, protonated ATP molecules exhibit diminished electrostatic repulsion and consequently promote condensation in the presence of crowders. As pH increases, the deprotonation of ATP molecules enhances their electrostatic repulsion and therefore inhibits condensate formation, accompanied by a decreased turbidity. The recovered turbidity at high pH indicates the reentrant formation of ATP condensates which is potentially caused by increased concentration of Na^+^ ions. For example, at pH 10, the Na^+^ ion concentration is approximately 160 mM, which is sufficient in screening electrostatic repulsion among ATP molecules and facilitates their condensation (Fig. 2C). As the ATP concentration is diluted and moves toward the binodal curve, this reentrant condensation becomes more pronounced. In particular, as pH increases from 3.26 to 11.64, ATP condensates form in acidic environments, dissolve at neutral pH, and then recover in basic environments (Fig. 4D and fig. S20B).

Moreover, ATP condensates are very sensitive to concentration variations. A twofold dilution of ATP condensates triggers their rapid dissolution (Fig. 4E and fig. S21), where ATP condensates exhibit gradient-directed migration and swim to their dissolution, driven by a dialytaxis mechanism (movie S1) (*53*). Subsequently, water evaporation at 50 °C leads to a gradual recovery of ATP condensates (Fig. 4F, fig. S21, and movie S2). This process begins with stochastic nucleation of micro-condensates, followed by their spontaneous growth and coalescence into larger droplets upon contact (Fig. 4F and movie S2). These dynamic and adaptive behaviors of ATP condensates are reminiscent of some biomolecular condensates in cellular spaces that dynamically assemble, dissolve, and adapt their composition in response to environmental fluctuations (*54-57*). The environment-responsive properties of ATP condensates also represent a dynamic form of compartmentalization with potentials as drug-carriers or prebiotic protocells.

### Selective and dynamic molecular partitioning within ATP condensates

Beyond their responsive properties, ATP condensates further show their capabilities to selectively take up guest molecules (Fig. 4G). Small-molecule fluorophores, such as propidium iodide and calcein, preferentially partition into ATP condensates (Fig. 4H). Similarly, oligonucleotides, oligopeptides, and macromolecules (e.g., polypeptides and DEX) accumulate within ATP condensates (Fig. 4H and fig. S22). In contrast, hydrophobic molecules, such as ATTO 647, Nile Red, and rhodamine B, are excluded outside from condensates (Fig. 4H and fig. S22). These results indicate that ATP condensates could enrich relatively hydrophilic guest molecules, while the continuous phase preferentially accommodates more hydrophobic guest molecules. Such difference between ATP condensates and the continuous phase enables the selective partitioning of guest molecules based on their physicochemical properties.

In addition to selective partitioning, we further demonstrate that ATP condensates can dynamically encapsulate and release molecular cargos triggered by external stimuli. Using calcein as an example, we observe a reversible temperature-gated sequestration within and release from ATP condensates (Fig. 4I and movie S3). In particular, the partition coefficient of calcein increases as ATP condensates form when the temperature exceeds the LCST and returns to 1 with the dissolution of condensates upon cooling for multiple cycles (Fig. 4I). The ability of ATP condensates to reversibly sequester guest molecules resembles biological condensates that dynamically regulate molecular transport and distribution to dictate biochemical processes (*58-60*).

### Protection of RNA cleavage in ATP condensates

Having demonstrated the ability of ATP condensates to concentrate biomolecules, we further explore how ATP condensates may regulate enzymatic reactions. To this end, we select an RNA cleavage reaction by DNAzyme as a model system given its importance in RNA processing and prebiotic chemistry (Fig. 5A) (*61, 62*). The RNA substrate is labeled with a FAM fluorophore at the 5’ terminus and a BHQ quencher at the 3’ terminus, exhibiting minimal fluorescence when intact due to proximity quenching and enabling real-time kinetic measurement of cleavage based on fluorescence increment. Fluorescence microscopy shows that the DNAzyme (E), RNA substrate (S), and cleaved fragments (S1 and S2) all preferentially partition into ATP condensates with over 10-fold local enrichment (Fig. 5, B and C).

**Fig. 5.**
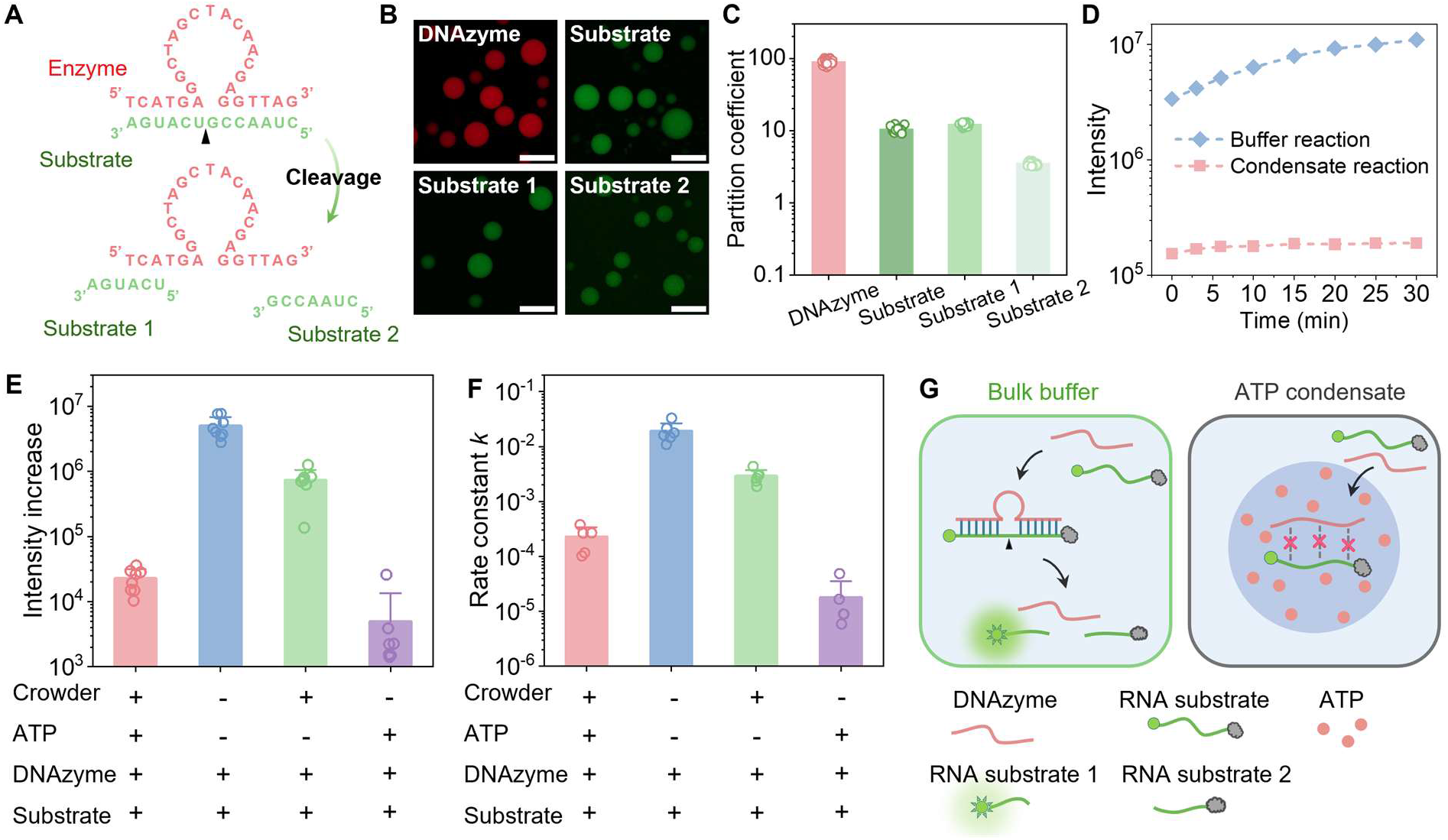
RNA is protected from cleavage in ATP condensates. (**A**) Schematic illustration of nucleotide sequences and the cleavage of RNA substrate by a DNA enzyme into two RNA segments, substrate 1 and substrate 2. RNA substrates were labeled with fluorophores and quenchers at their 5’ and 3’ ends. The cleavage dissociates the fluorophore from the quencher, enabling fluorescence emission. (**B**) Representative fluorescence images and (**C**) partition coefficients of DNAzyme (2.5 µM), RNA substrate (2 µM), cleaved RNA substrate 1 (2 µM) and substrate 2 (2 µM) within ATP condensates. Scale bars are 50 μm. (**D**) Time-lapsed fluorescence intensity profiles of RNA cleavage reaction in ATP condensate and buffer solutions. Comparison of (**E**) the fluorescence increment and (**F**) the reaction rate constant *k* of RNA cleavage reaction under various conditions. The rate constants are obtained by fitting experimental data to single exponential growth. For each condition, at least five independent experiments are performed to ensure statistical reliability. (**G**) Schematic illustration of RNA cleavage reaction in bulk buffers and suppressed RNA cleavage in ATP condensates.

Molecular enrichment and co-localization of enzyme and substrate normally increase reaction kinetics (*11, 63*). Surprisingly, despite this strong co-concentration, the RNA cleavage reaction is significantly suppressed within ATP condensates (Fig. 5D). Specifically, the fluorescence increment is reduced by over two orders of magnitude in ATP condensate solution (Fig. 5E, red column) compared to bulk buffer (Fig. 5E, blue column). Fitting fluorescence kinetics to an exponential model reveals an 84-fold reduction in reaction rate from 1.90×10^−2^ ± 7.01×10^−3^/min in bulk buffer (Fig. 5F, blue column) to 2.26×10^−4^ ± 1.04×10^−4^/min in condensate solution (Fig. 5F, red column). Even when we increase the initial concentrations of enzyme and catalytic ions by tenfold, the reaction rate is still not restored, underscoring the profound RNA cleavage protection inside ATP condensates (fig. S23).

To understand this phenomenon, we first examine the effects of crowding on reactions. Firstly, the increased viscosity inside ATP condensates would proportionally reduce the diffusion coefficient of substrates and enzymes according to the Stokes-Einstein relation. However, as the viscosity has only increased by approximately 10 times (Fig. 1I), this reduced diffusion would be readily compensated by the increased local concentrations of enzymes and substrates (Fig. 5C). This is further supported by modestly reduced reactions in viscous PEG solutions (Fig. 5, E and F, green column). Therefore, it suggests that the protected RNA cleavage is not simply explained by a diffusion-limited model.

We then observe that ATP condensates establish an acidic internal environment with a pH of approximately 2.49 inside condensates, while pH is about 3.42 in the continuous phase (see methods for details). This acidic environment arises from ATP deprotonation, supported by a decreasing pH with increasing ATP concentration (fig. S24). To isolate the effect of pH on the reactions, we perform a series of cleavage reactions in buffers with varying pH in the absence of PEG and ATP. The reaction is indeed significantly suppressed in buffers of pH 3 compared to that in buffers of pH 7.5 (fig. S25). The suppression persists even when we increase the concentrations of substrate and enzyme by 10-fold (fig. S25). This acidic environment inside condensates may reduce the catalytic efficiency of enzymes and thus protect RNA from cleavage (*64, 65*).

Beyond the pH effect, we find that ATP molecules alone could also significantly suppress the cleavage reaction, with both fluorescence increment and reaction rate reduced by three orders of magnitude compared to bulk buffer (Fig. 5, E and F, purple column). These reactions are restricted in the presence of ATP even at a neutral pH 7.5 (fig. S26). ATP molecules could potentially disturb the binding of enzymes to substrates and catalysts, leading to suppressed RNA cleavage reactions (*65-67*). Collectively, our results demonstrate that ATP condensates could protect RNA from cleavage by creating an acidic microenvironment and potentially interfering with the substrate-enzyme interaction (Fig. 5G). These results indicate that ATP condensates could function as reservoirs to store and protect RNA against cleavage and degradation, underscoring a dual role of ATP as structural building blocks for condensate formation as well as biochemical regulators on localized enzymatic reactions.

## Conclusion

This work reveals an important yet previously unrecognized role of ATP in forming protein-free condensates via homotypic interactions in crowded environments. Mechanistically, screened electrostatic repulsion and enhanced hydrogen bonding overcome the enthalpic and entropic barriers of ATP, enabling the formation of liquid-like ATP condensates in segregation with crowders. This discovery expands the paradigm of ATP-enriched condensates beyond the long-standing focus on associative ATP-polycation condensates. Importantly, ATP condensates exhibit distinct thermodynamic properties compared to typical ATP-polycation condensates formed by associative LLPS. For example, ATP condensates exhibit high fluidity and dynamic responsiveness to environmental changes, in contrast to very stable ATP-polycation condensates reported in previous studies (*19, 20, 31, 51*). Such dynamic responsiveness further enables ATP condensate to reversibly concentrate and release guest molecules. Beyond that, ATP condensates create unique chemical microenvironments to sequester ribonucleic acids and protect them from cleavage. Given the critical roles of ATP in biology, the formation mechanism and environment-responsive properties of ATP condensates potentially provide insights into prebiotic chemistry, where small-molecule LLPS enables dynamic compartmentalization, adaptation, and protection of prebiotic molecules against environmental stress (*68-70*).

## Acknowledgments

We thank Prof. Tuomas Knowles and Prof. Boon Leong Lim for helpful discussions and comments on this project.

## Funding

This work was supported by the General Research Fund (Nos. 17306221, and 17317322) and Collaborative Research Fund (C7165-20GF) from the Research Grants Council (RGC) of Hong Kong, the National Natural Science Foundation of China (NSFC)-RGC Joint Research Scheme (N_HKU718/19), as well as NSFC (No. 22525805). H.C.S. was funded in part by the RGC Senior Research Fellow (SRFS2425-7S04) by RGC and the Health@InnoHK program of the Innovation and Technology Commission of the Hong Kong SAR Government.

## Author contributions

F.C. conceived the research; F.C. and Y.W. designed the research and experiments; Y.W., F.C., and P.D.W. performed the experiments and analyzed data; F.C. and H.C.S. supervised the research. Y.W. and F.C. wrote the paper, and all authors have revised it.

## Competing interests

Ho Cheung Shum is a scientific advisor of EN Technology Limited, MicroDiagnostics Limited, PharmaEase Tech Limited, Upgrade Biopolymers Limited and Multera Limited, in which he owns some equity, and is a founding director and co-director of the research center, Advanced Biomedical Instrumentation Centre Limited. The works in this paper are, however, not directly related to the works of these entities, as far as we know. The authors declare no other competing interests.

## Data, code, and materials availability

All data are available in the main text or the supplementary materials, and also from the corresponding authors upon request.

