## Supplementary Material for "Adenosine 5’-triphosphate (ATP) forms protein-free and responsive condensates in crowded environments"

**The PDF file includes:**

Materials and Methods
Figs. S1 to S27

**Other Supplementary Materials for this manuscript include the following:**

Movies S1 to S3

**Materials and Methods**

Materials

Adenosine 5'-triphosphate (ATP), Adenosine 5'-diphosphate (ADP), Guanosine 5'-triphosphate (GTP), Cytidine 5'-triphosphate (CTP), Poly(ethylene glycol) (PEG, Mw 8000), Fluorescein isothiocyanate labeled dextran 10,000 (FITC-Dex 10k), Fluorescein isothiocyanate labeled dextran 70,000 (FITC-Dex 70k), Poly-L-lysine hydrobromide (PLL, Mw~30,000-70,000), Fluorescein isothiocyanate labeled Poly-L-lysine (FITC PLL, Mw~30,000-70,000), ethylene glycol, Poly(2-ethyl-2-oxazoline) (average Mw ~50,000), Ficoll 400, Urea, 1,6-Hexanediol, ATTO-647, Rhodamine B, Propidium Iodide, and Calcein were purchased from Sigma-Aldrich. Potassium Chloride (2M solution), Sodium Chloride (5M solution), Magnesium Chloride (1M solution), and Nile red were purchased from Thermo Fisher Scientific. Calcium Chloride (1M solution) and Dextran 10 (Mw ~10,000) were purchased from Macklin. Poly(ethylene glycol) (PEG, average Mw ~2000 and 600), Rhodamine-labeled PEG (Rh-PEG, Mw ~10,000) and Fluorescein isothiocyanate(FITC) were purchased from Aladdin. Fluorescent polystyrene microspheres (size: 1 μm, concentration: 10mg/ml, excitation peak: 488 nm) was purchased from Tianjin Junvijia Technology Co. Ltd. Cy5-labeled Single-stranded RNA (Cy5-UUAGAAUUAGAAUUAGAAUUAGAA), FAM-labeled Single-stranded DNA (FAM-TTAGAATTAGAATTAGAATTAGAA), Cy5-labeled and original DNAzyme (TCATGAGGCTAGCTACAACGAGGTTAG), FAM and BHQ-labeled RNA substrate (CUAACCGUCAUGA), FAM-labeled cleaved substrate fragments S1 (CUAACCG) and S2 (UCAUGA) were synthesized by IDT. FAM-labeled R10 (5,6-FAM-RRRRRRRRRR) was synthesized by CASLO. Milli-Q water (18.2 MΩ, pH=7) was used in all experiments. All chemicals were used as received without further purification.

Condensate formation and sample preparation

Stock solutions of nucleotides were firstly prepared at high concentrations (500mM) using Milli-Q water (18.2 MΩ, pH=7) and stored under -20 °C. Stock solutions of crowders were firstly prepared at high concentrations (500mg/mL) using Milli-Q water (18.2 MΩ, pH=7) and stored under room temperature. Solutions were mixed and vortexed thoroughly to form condensates based on final solute concentrations. Samples contain 350 mg/ml crowders and 80mM nucleotides are used unless otherwise noted. Samples were added into chambers where spacers with 9 mm diameter and 0.12 mm deep are sandwiched by cover slides for imaging and observation.

Characterization of ATP and crowder phases

Sample solutions with 350 mg/ml PEG and 80mM ATP were prepared in microcentrifuge tubes as stated in the condensate formation part and were centrifuged to stratify into two immiscible phases. The PEG-enriched phase was distinguished by taking 1 uL of sample from each of the two phases and mixing with 1 uL of 160 mM Dextran solutions (Mw 10,000). The ATP-enriched phase was distinguished by taking 0.5 uL of sample from each of the two phases and mixing with 1 uL of 20 mM PLL solution and diluting with water to varying extents.

Fluorescence recovery after photobleaching (FRAP)

FRAP experiments were conducted using a Carl Zeiss LSM 980 confocal laser scanning microscope, equipped with a 40× dry objective. Samples were prepared at a final concentration of 80 mM ATP and 350 mg/mL PEG 8K with 5 μM FITC-labeled PLL (Mw 30-70k). A circular region of interest was bleached with a 488-nm laser at 100% power for 4 seconds and subsequent recovery of the bleached area was recorded with a 488-nm laser. For analysis, FRAP recovery trace was plotted and fitted by software ImageJ.

Single-particle tracking

Single-particle tracking was based on the images extracted from video recordings. Particle trajectories were reconstructed in two dimensions by extracting the x-y coordinates of individual particles using custom MATLAB scripts. The mean squared displacement (MSD) for each trajectory was calculated by formula $\mathrm{MSD}(\Delta t) = \langle (x(t+\Delta t) - x(t))^2 \rangle + \langle (y(t+\Delta t) - y(t))^2 \rangle$, where $x$ and $y$ represent the particle coordinates and $\Delta t$ denotes the lag time. The two-dimensional diffusion coefficient ($D$) was derived by fitting MSD curves with $\mathrm{MSD}(\Delta t) = 4D\Delta t^{\alpha}$, where $\alpha$ is the scaling exponent and $D$ is the diffusion coefficient. The internal viscosity ($\eta$) of condensates was calculated based on the Stokes-Einstein equation $\eta = \frac{k_B T}{6\pi D R}$, where $k_B$ is the Boltzmann constant, $T$ is the absolute temperature, and $R$ is the particle radius.

Turbidity measurement

Turbidity of ATP condensates was measured using a microplate reader (Molecular Devices). Solutions were prepared in 96-well plates with a total volume of 200 µL per well. Absorbance was monitored at a fixed wavelength of 500 nm. Following background correction, turbidity (%) was calculated by normalizing the corrected absorbance to the maximum absorbance value.

Experimental measurement of phase diagram

To construct the phase diagram of the PEG-ATP system, we employed turbidity tests coupled with optical microscopy. Stock solutions of ATP and PEG at high concentrations were first mixed at different initial ratios. Each mixture was then subjected to stepwise dilution which generated a comprehensive series of data points across a wide range of concentrations and composition ratios. The phase behavior of each point was determined by the solution turbidity. Samples near the phase boundary, marking the onset of phase separation, were visually examined the presence of condensates by microscopic confirmation. The phase diagram was plotted by the data points on a two-dimensional concentration diagram and the binodal curve, indicating the boundary between the single-phase and phase separation regions, was obtained by fitting these experimentally determined points.

NMR spectroscopy

$^1$H NMR spectra were acquired using a Bruker Avance III 500 MHz spectrometer equipped with a Prodigy BBO cryoprobe at 298.15 K. NMR samples were prepared in solvents ($H_2O$ : $D_2O$ = 9 : 1) and measured using a water-suppression pulse with 64 scans. Chemical shifts were determined in reference to the solvent residual peaks. All data was processed in MestReNova software.

Temperature effects

ATP condensates were prepared at room temperatures and added into observation chambers mounted on a transparent thermal plate (Tokai Hit Ltd.). The target temperatures were set by a TPi controller and the real temperature was monitored by a temperature sensor. Vaseline was used to seal the samples once they are loaded into chambers to minimize solvent changes caused by temperature fluctuations. Samples were observed following a 5-minute equilibration period after reaching the target temperature. The partition coefficient evolution in ATP-PEG solution ($c_{\mathrm{PEG}}$ = 175 mg/mL and $c_{\mathrm{ATP}}$ = 175mM) in response to temperature changes was characterized by Calcein with a final concentration of 0.25mg/mL. The fluorescent intensity of three thermal cycles was measured and obtained by a Carl Zeiss LSM 980 confocal laser scanning microscope.

Concentration responsiveness

For dilution-induced condensate dissolution, 5 uL of water was added to dilute 5 uL of ATP condensate solution (350 mg/mL PEG, 80 mM ATP). The condensation process was achieved by evaporating 10 uL ATP-PEG solution (150 mg/mL PEG, 40 mM ATP) 50 °C. The dissolution and condensation process were observed with a Leica inverted microscope.

Guest molecule partitioning

Fluorescent guest molecules were added to the solutions before condensates formed. ATP condensates are formed with the final solution of 350 mg/mL PEG 8K and 80 mM ATP. The fluorescent images were captured by a Carl Zeiss LSM 980 confocal laser scanning microscope. The final dye concentrations are ATTO 647 (25 μM), Rhodamine B (0.5mg/mL), Nile Red (0.05mg/mL), PI (0.5 μM), Calcein (0.25mg/mL), Cy5-labeled RNA (Cy5-UUAGAAUUAGAAUUAGAAUUAGAA, 2.5 μM), FAM-labeled DNA (FAM-TTAGAATTAGAATTAGAATTAGAA, 2.5 μM), FAM-labeled R10 ((5,6-FAM)-RRRRRRRRRR, 1 μM), FITC-labeled PLL (Mw 30-70k, 1 μM), respectively. Partition coefficients are measured as the fluorescence intensity ratio between condensate interior and continuous phase.

DNAzyme cleavage reaction

The DNAzyme cleavage reaction was incorporated into different buffer and condensate solutions. DNAzyme stock solution was heated at 95 °C for 3 min and cooled down at room temperature for 10 min. The DNAzyme was then added to different solution groups containing 1mM $MgCl_2$ and 40mM Tris-HCl buffer to achieve a final concentration of 1 μM. The reaction was initiated by the addition and mixing of RNA substrate stock solution at a final concentration of 0.5 μM. 100 μL reaction solutions were loaded into 96 well plates immediately and the cleavage reactions were monitored by FAM fluorescence intensity measured over time by a microplate reader (Molecular Devices) at excitation wavelength 485 nm and detection wavelength 525 nm. The regulatory effects on reaction were compared based on fluorescence intensity increase and reaction rate constant for 30 minutes. The fluorescence intensity increase was calculated by subtracting the starting fluorescence from the maximum fluorescence. The reaction rate constant was obtained by fitting the reaction kinetic profiles to single exponential equation $I(t) = 1 - e^{-kt}$, where $I(t)$ is the normalized fluorescence intensity, $t$ is the reaction time and $k$ is the rate constant.

pH measurement

The pH within ATP condensates was determined using pH-sensitive FITC as probe. A calibration curve for FITC probe was constructed by calibration solutions with 2 μM FITC at a range of pH values (fig. S27). Calibration solutions were prepared in 40 mM Tris buffer and the pH values were adjusted through the addition of HCl and NaOH (measured with Oakton pH 550 pH meter). Solutions were loaded in 96-well plates with a total volume of 100 μL per well and the fluorescence intensity was measured by a microplate reader (Molecular Devices) at excitation wavelength 485 nm and detection wavelength 525 nm. To measure the pH of condensates and continuous phase, samples were prepared as described above and the condensate phase and continuous phase were separated via centrifugation at 9000 rpm for 5 minutes. Separated solutions were dispensed in equal volume and added with FITC to a final concentration of 2 μM for fluorescence measurement. The pH was extrapolated from the calibration curve and validated using both calibrated pH meter and pH test strips, both of which showed consistent results for condensate phase. However, the FITC-based measurement of the PEG-rich continuous phase was unreliable, likely due to altered photophysical properties of FITC and strong microenvironment effects in the extremely crowded PEG solution. Therefore, the pH of the continuous phase was directly measured using a pH meter, with additional validation via pH test strips.

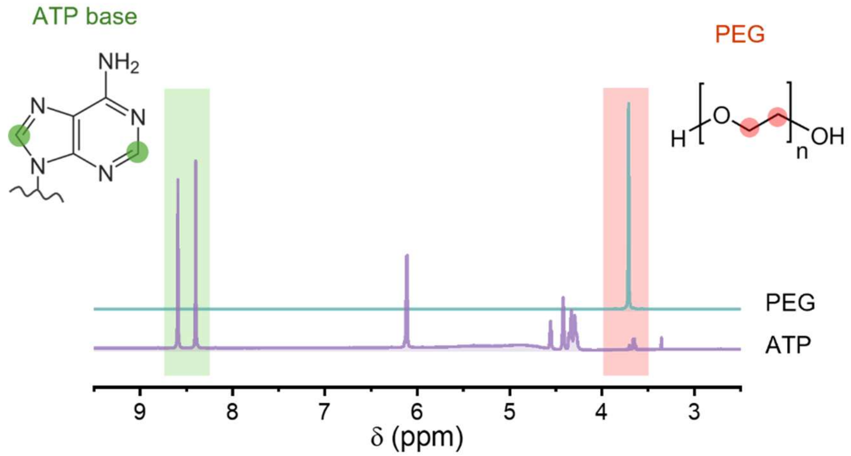


**Fig. S1.**

A typical $^{1}$H NMR spectrum of PEG protons (top) and ATP adenine base protons (bottom) acquired with 25 mg/mL PEG and 25 mM ATP respectively.

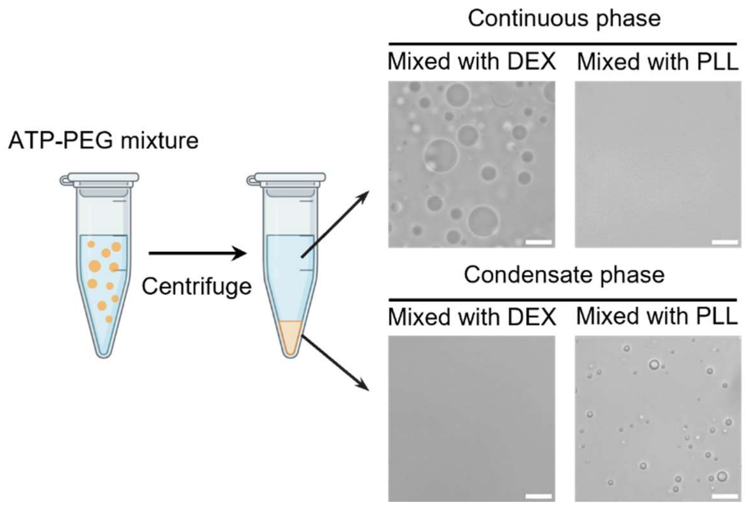


**Fig. S2.**

Composition determination of the condensates and the continuous phase ($c_{PEG}$= 350 mg/mL and $c_{ATP}$= 80 mM). Following centrifugation, condensates accumulate at the bottom, while the continuous phase resides as the top layer. The two immiscible phases are extracted and mixed with dextran and PLL, respectively. Scale bars are 10 μm.

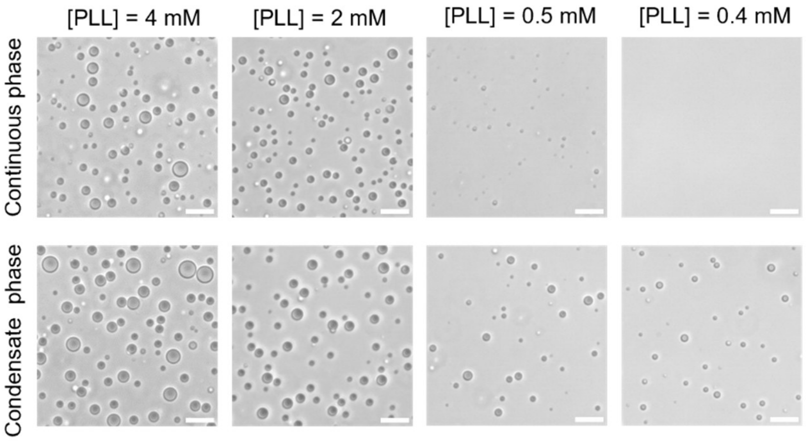


**Fig. S3.**

Representative images of 0.5 μL continuous phase (Top) and condensate phase (Bottom) of centrifuged ATP-PEG solution ($c_{PEG}$= 350 mg/mL and $c_{ATP}$= 80 mM) mixed with 1 μL 20 mM Poly-L-lysine (PLL, 30-70 kDa) and diluted to different extents. Scale bars are 10 μm.

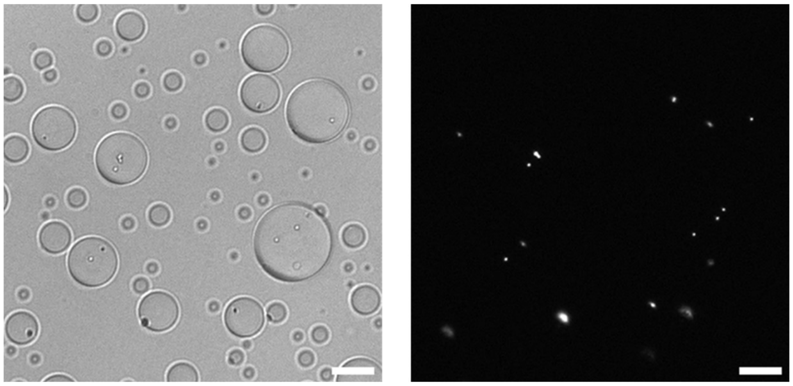

**Fig. S4.**
Bright field and fluorescence images of ATP condensates with fluorescent microparticles partitioned inside. Scale bars are 20 µm.

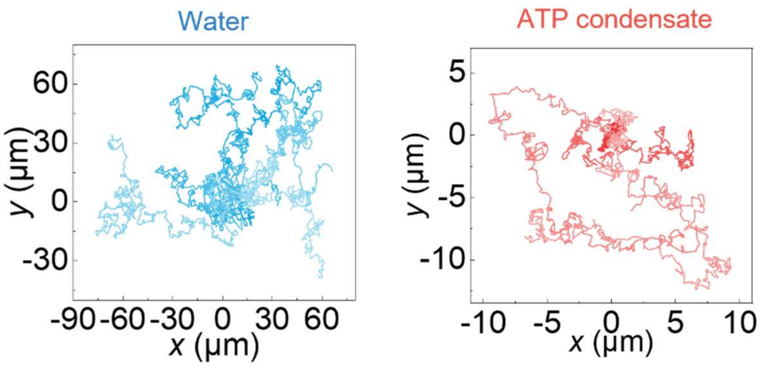


**Fig. S5.**
2D trajectories of fluorescent microparticles in water and ATP condensates. Different colors in each plot represent independent tracked particles ($n$ = 7).

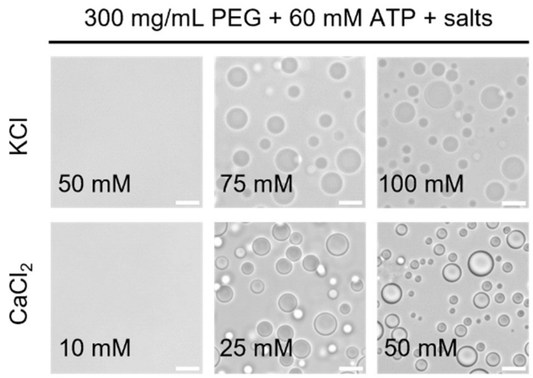


**Fig. S6.**

Representative images of mixtures of 300 mg/mL PEG and 60 mM ATP under different additive concentrations. Scale bars are 10 μm.

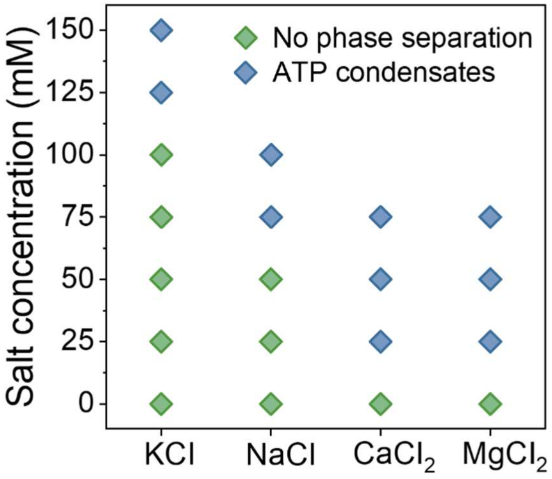


**Fig. S7.**

Schematic representation of the critical salt concentration for condensate formation from ATP solution ($c_{\mathrm{PEG}}$= 300 mg/mL, $c_{\mathrm{ATP}}$= 50 mM) with different added salts.

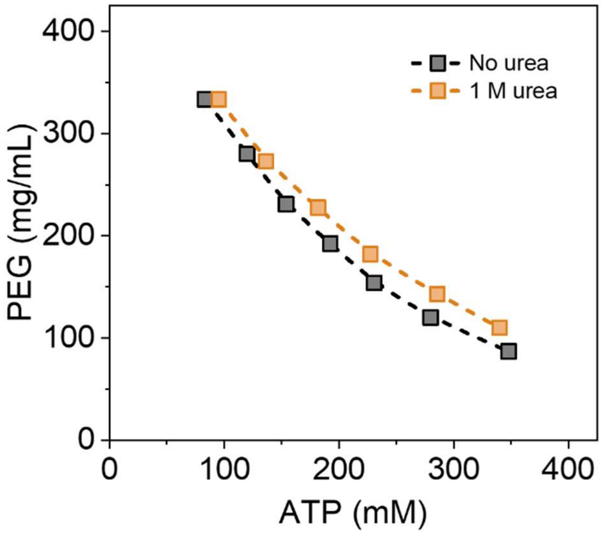


**Fig. S8.**
Phase diagrams and binodal curves of ATP−PEG mixture with and without the addition of 1 M urea.

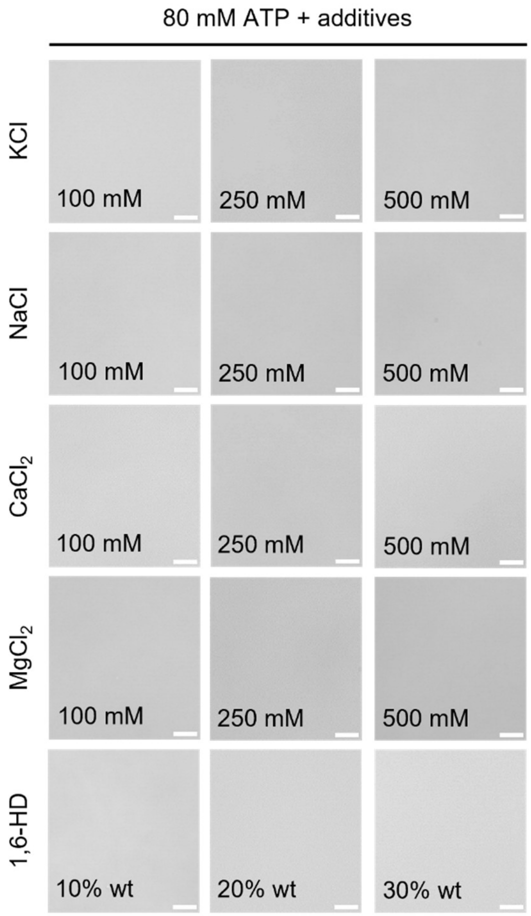


**Fig. S9.**

Representative images of 80 mM ATP only under different additive concentrations. Scale bars are 10 μm.

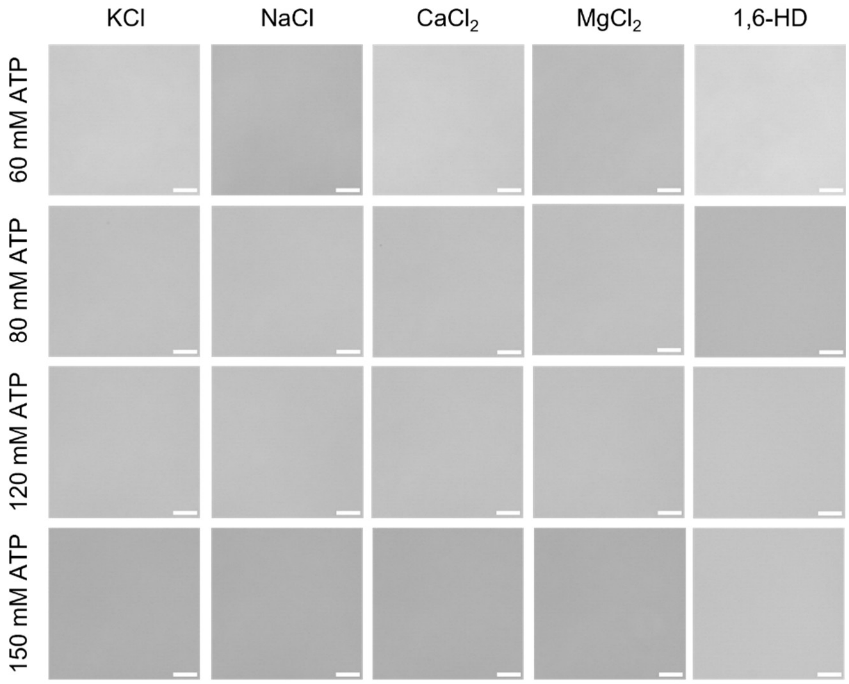


**Fig. S10.**

Representative images of only different concentrations of ATP with various kinds of additives (250 mM salts and 20% wt 1,6-HD). Scale bars are 10 μm.

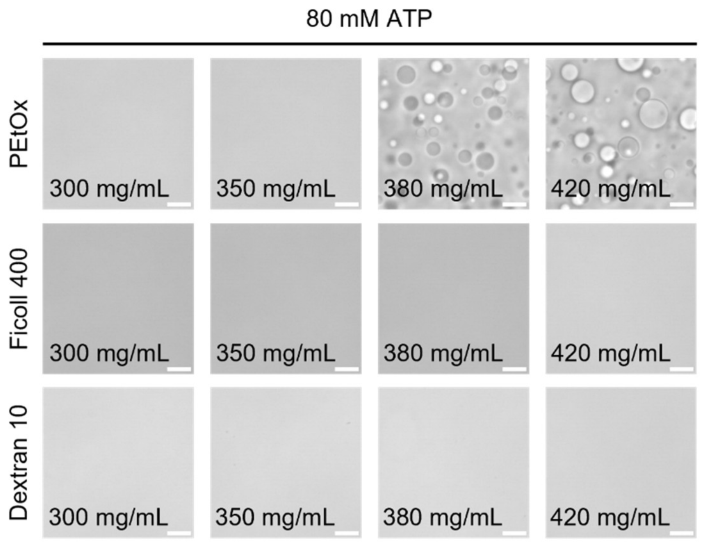


**Fig. S11.**

Representative images of ATP solutions added with different crowders with constant ATP concentration of 80 mM. Scale bars are 10 μm.

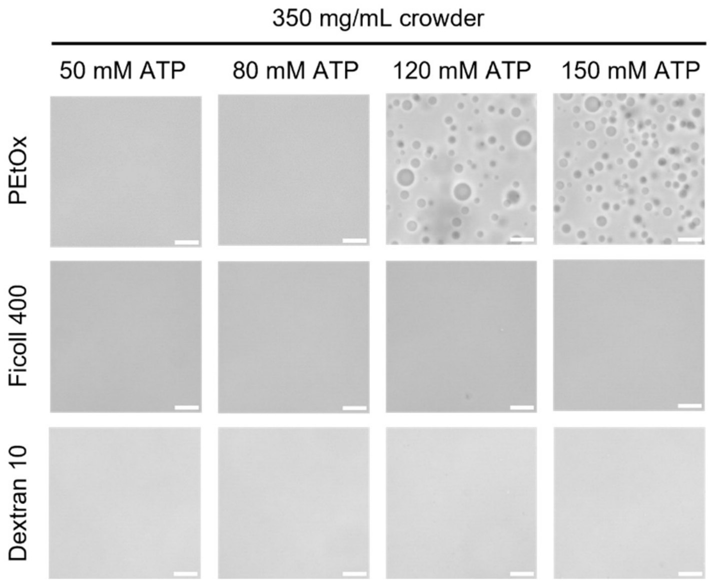


**Fig. S12.**

Representative images of ATP solutions added with different crowders with constant crowder concentration of 350 mg/mL. Scale bars are 10 μm.

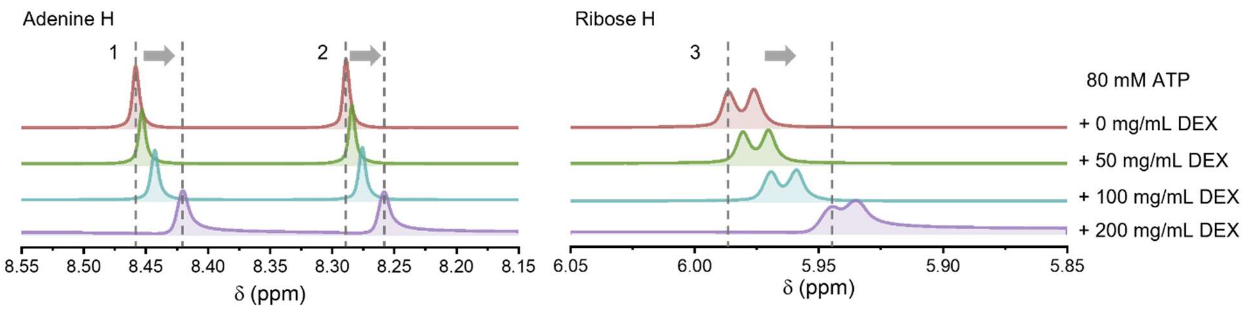


**Fig. S13.**

A typical $^{1}$H NMR spectrum of ATP adenine base and ribose sugar protons acquired with 80 mM ATP and increasing DEX concentrations (from top to bottom).

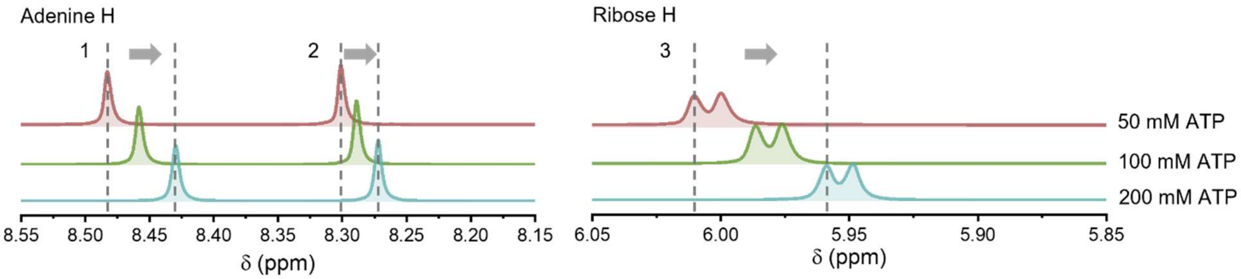


**Fig. S14.**

A typical [1]H NMR spectrum of ATP adenine base and ribose sugar protons acquired with increasing ATP concentrations (from top to bottom).

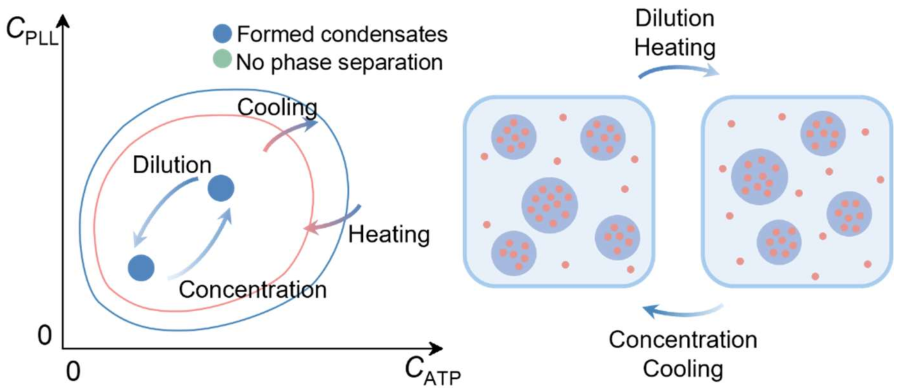


**Fig. S15.**

Schematic illustration of the stability of associative ATP-PLL condensates.

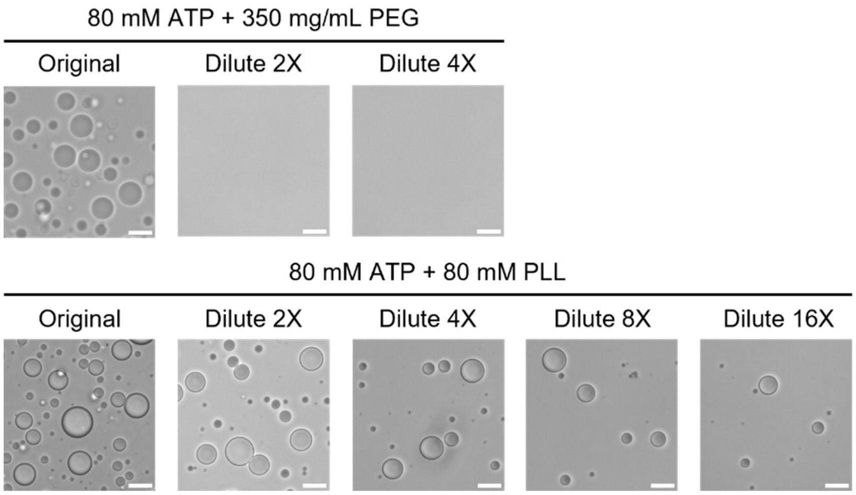


**Fig. S16.**

Comparison between segregative ATP condensates and associative ATP-PLL condensates by dilution. Scale bars are 10 μm.

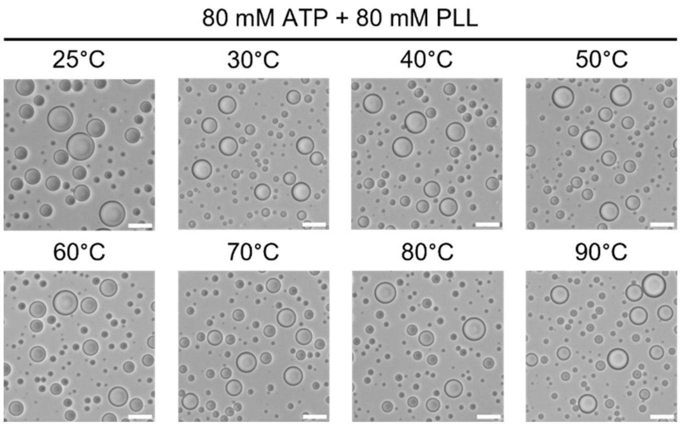


**Fig. S17.**
Thermal stability test of associative ATP-PLL condensates. Scale bars are 10 μm.

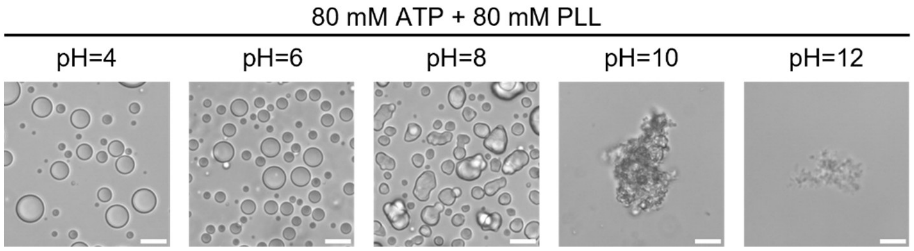


**Fig. S18.**

Representative images of associative ATP-PLL condensates under different pH. Scale bars are 10 μm.

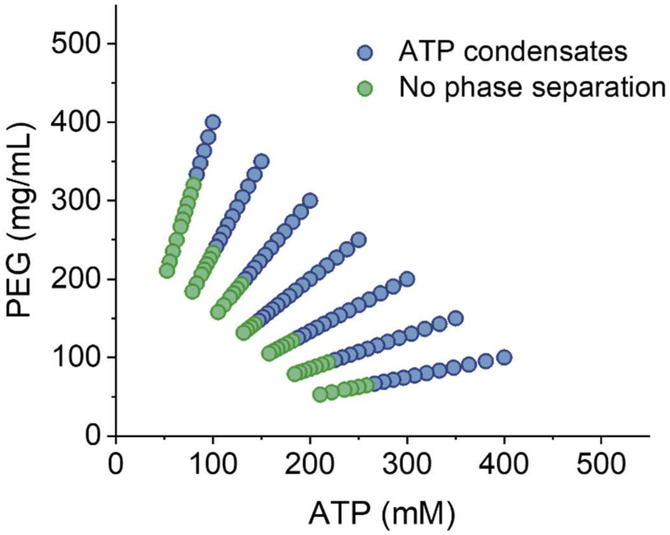


**Fig. S19.**
Phase diagram of ATP-PEG mixture at 60 °C.

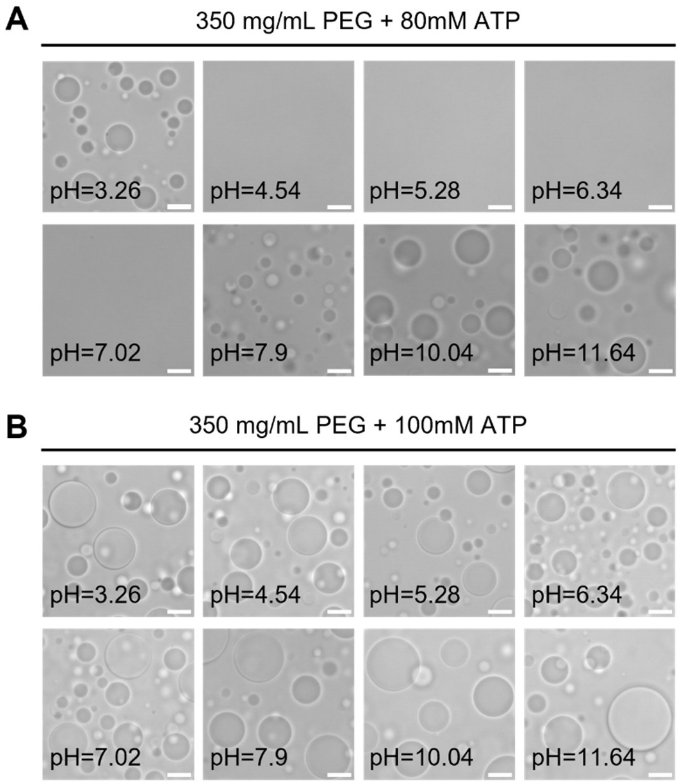


**Fig. S20.**

Representative images of (A) ATP-PEG solutions ($c_{PEG}$= 350 mg/mL and $c_{ATP}$= 80 mM) and (B) ATP-PEG solutions ($c_{PEG}$= 350 mg/mL and $c_{ATP}$= 100 mM) under different pH. Scale bars are 10 μm.

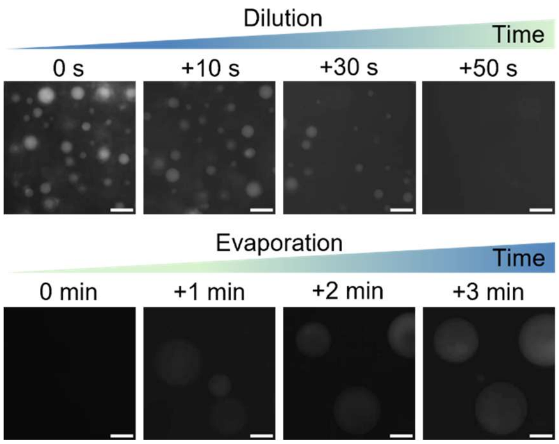


**Fig. S21.**

Fluorescence images of the dissolution of ATP condensates ($c_{PEG}$= 350 mg/mL and $c_{ATP}$= 80 mM) upon twofold dilution with water and their subsequent reformation via water evaporation induced by heating at 50 °C. Scale bars, 100 µm.

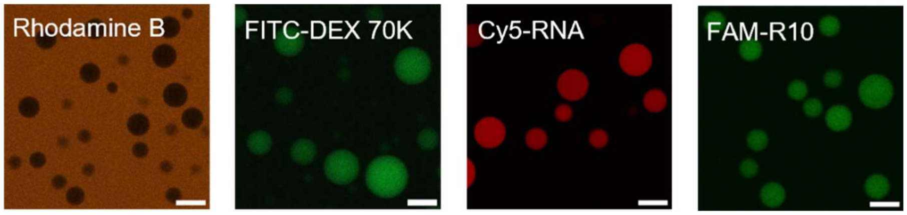


**Fig. S22.**

Fluorescence image showing the selective partitioning of diverse guest molecules in ATP condensates ($c_{PEG}$= 350 mg/mL and $c_{ATP}$= 80 mM) or in the continuous phase. Scale bars are 20 µm.

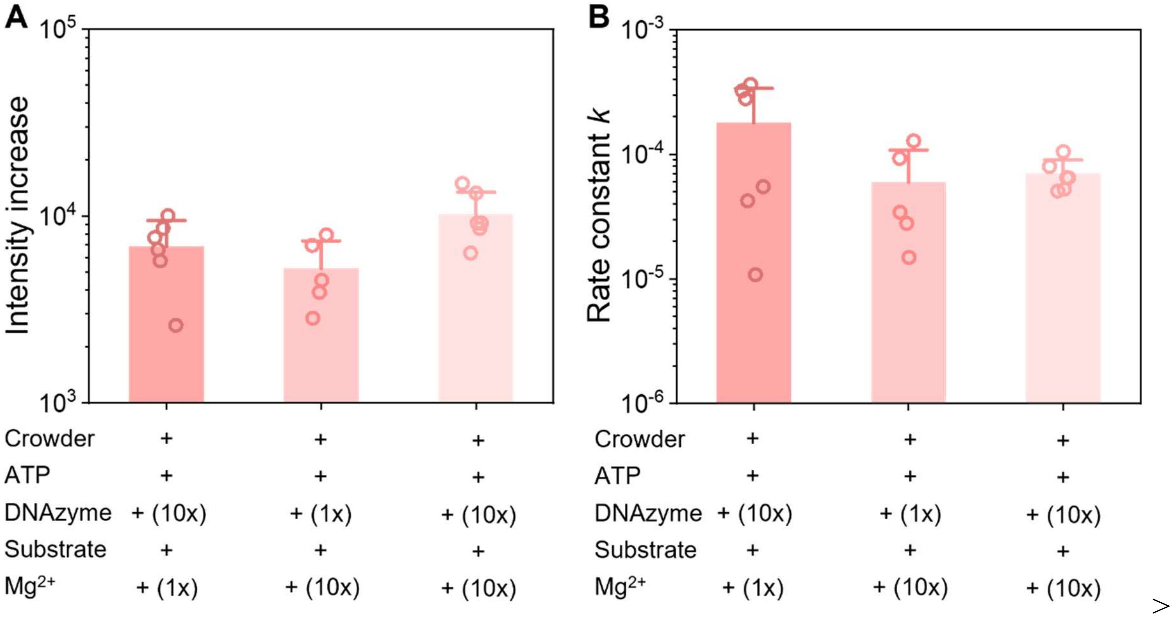


**Fig. S23.**

Comparison of (A) the fluorescence increment and (B) the reaction rate constant *k* of RNA cleavage reactions in ATP condensate system under various conditions.

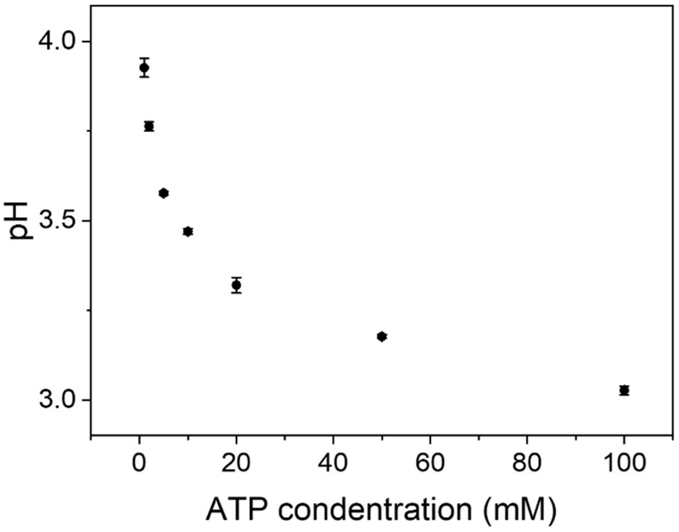


**Fig. S24.**
Measured pH values of ATP solution with different concentrations ($n$ = 3).

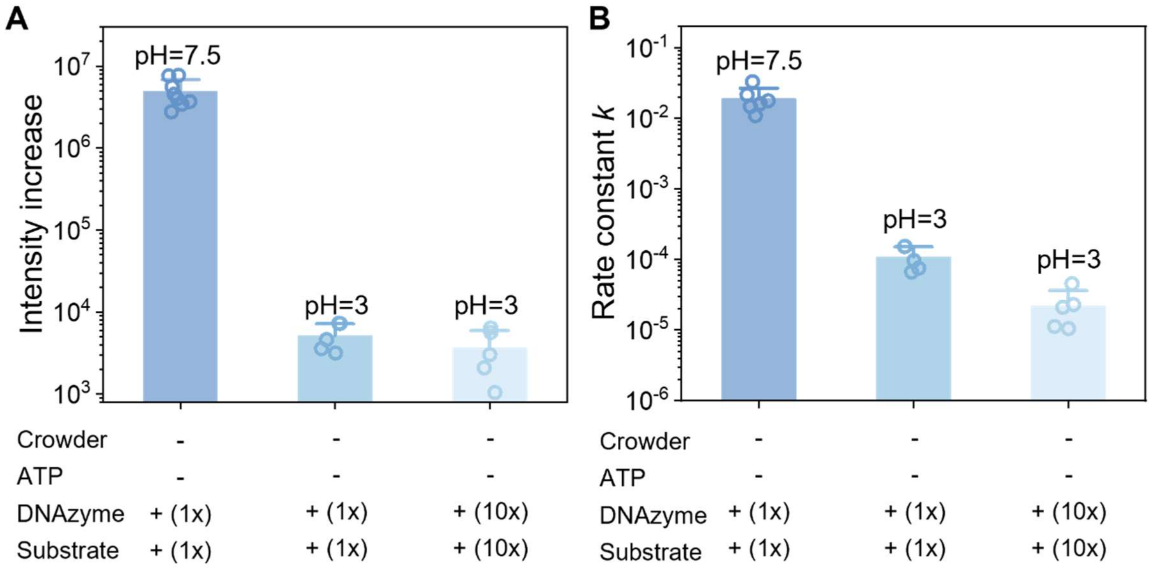


**Fig. S25.**

Comparison of (A) the fluorescence increment and (B) the reaction rate constant *k* of RNA cleavage reactions in buffer system with pH=3 (the blue and green columns) and pH=7.5 (the red column).

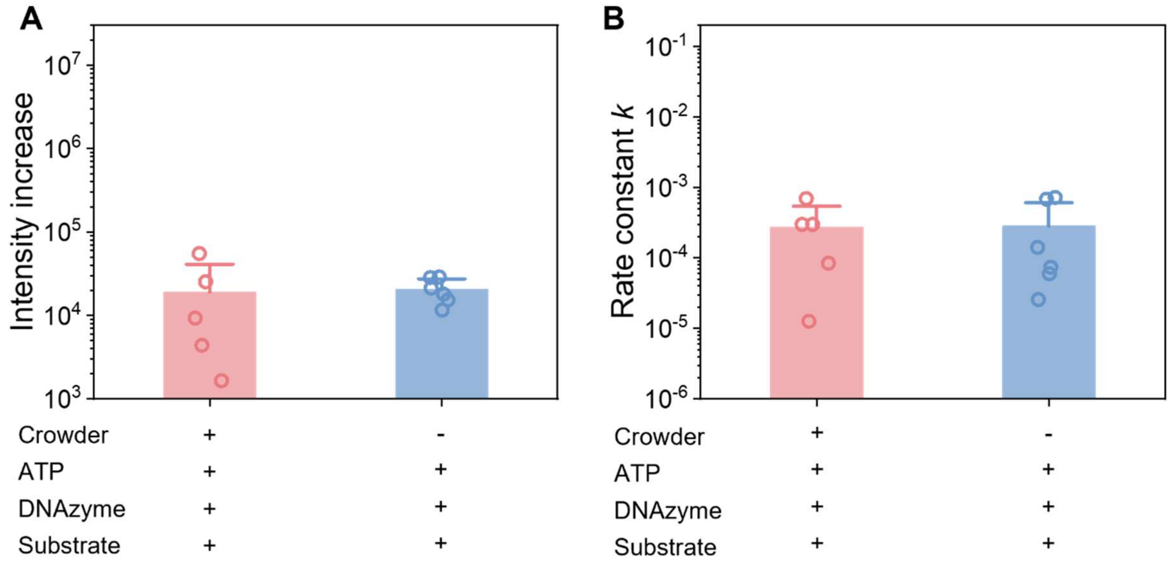


**Fig. S26.**

Comparison of (A) the fluorescence increment and (B) the reaction rate constant $k$ of RNA cleavage reactions in ATP condensate solution with pH=7.5 (the red column) and pure ATP solution (the blue column) with pH=7.5.

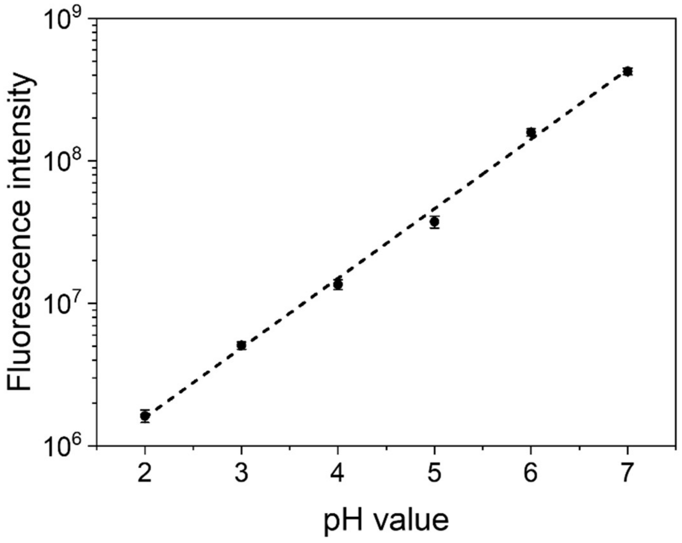


**Fig. S27.**

Calibration curve (linear fit with $R^2$ of 0.99775) for FITC probe at varying pH conditions prepared in 40 mM Tris aqueous buffer. pH condition is adjusted through the addition of HCl and NaOH ($n$ = 4).

**Movie S1.**
Dissolution of ATP condensates upon twofold dilution with water.

**Movie S2.**
Reformation of ATP condensates via water evaporation induced by heating at 50 °C.

**Movie S3.**
The partitioning of Calcein into ATP condensates controlled by temperature.
